# Genomic Epidemiology of *Escherichia coli* Isolates from a Tertiary Referral Center in Lilongwe, Malawi

**DOI:** 10.1101/2020.05.22.106062

**Authors:** Gerald Tegha, Emily J. Ciccone, Robert Krysiak, James Kaphatika, Tarsizio Chikaonda, Isaac Ndhlovu, David van Duin, Irving Hoffman, Jonathan J. Juliano, Jeremy Wang

## Abstract

Antimicrobial resistance (AMR) is a global threat, including in sub-Saharan Africa. However, little is known about the genetics of resistant bacteria in the region. In Malawi, there is growing concern about increasing rates of antimicrobial resistance to most empirically used antimicrobials. The highly drug resistant *Escherichia coli* sequence type (ST) 131, which is associated with the extended spectrum β-lactamase *bla_CTX-M-15_*, has been increasing in prevalence globally. Previous data from isolates collected between 2006-2013 in southern Malawi have shown the presence of ST131 and the *bla_CTX-M-15_* gene in the country. We performed whole genome sequencing (WGS) of 58 clinical *E. coli* isolates at Kamuzu Central Hospital, a tertiary care center in central Malawi, collected from 2012-2018. We used Oxford Nanopore Technologies (ONT) sequencing, which was performed in Malawi. We show that ST131 has become more prevalent (14.9% increasing to 32.8%) and that the *bla_CTX-M-15_*gene is occurring at a higher frequency (21.3% increasing to 44.8%). Phylogenetics show isolates are highly related between the central and southern geographic regions and confirm that ST131 isolates are contained in a single group consistent with recent expansion. All AMR genes, including *bla_CTX-M-15_*, were widely distributed across sequence types. We also identified an increased number of ST410 isolates, which in this study tend to carry a plasmid-located copy of *bla_CTX-M-15_* gene at a higher frequency than *bla_CTX-M-15_* occurs in ST131. This study confirms the expanding nature of ST131 and the wide distribution of the *bla_CTX-M-15_* gene in Malawi. We also highlight the feasibility of conducting longitudinal genomic epidemiology studies of important bacteria with the sequencing done on site using a nanopore platform that requires minimal infrastructure.

**DATA SUMMARY:** The sequencing data used for this analysis is available in public data repositories. Information on the sequences used is provided in Supplementary Table 2.

## INTRODUCTION

Antimicrobial resistance (AMR) is one of the most serious global public health threats.[1] Of specific concern are *Enterobacterales* (formerly the Enterobacteriaceae) that are resistant to third-generation cephalosporins such as ceftriaxone. The World Health Organization (WHO) has designated ceftriaxone-resistant *Enterobacterales* as a critical priority.[2] In sub-Saharan Africa (SSA), there is growing evidence that ceftriaxone-resistant *Enterobacterales* are important pathogens in invasive infections such as bacteremia. Given that ceftriaxone is often used to treat severe infections in SSA, and carbapenems are not often available, this is a major concern.

Among the *Enterbacterales, Escherichia coli (E. coli)* is a common cause of invasive disease, accounting for between 3% and 33% of positive blood cultures in case series in Africa.[3–8] *E. coli* is becoming more resistant to commonly used antibiotics in SSA, including ceftriaxone. Additionally, recent evidence suggests that the highly drug resistant *E. coli* sequence type (ST) 131 has been increasing in prevalence globally.[9–11] This sequence type is an extraintestinal pathogenic *E. coli* (ExPEC) that is associated with bloodstream and urinary tract infections, often possessing genes associated with extended-spectrum β-lactamases (ESBL).[12, 13] The main mechanism of cephalosporin resistance is drug inactivation mediated by hydrolysis of the β-lactam ring by ESBL enzymes. CTX-M derivatives are the dominant and most widely distributed ESBL enzyme among *E. coli*.[14] CTX-M-15 is strongly associated with ST131.[15–18] The global spread of ESBL-*E. coli* is largely attributed to the dissemination of *E. coli* strains carrying the *bla_CTX-M-15_* gene, especially *E. coli* O25b:H4-ST131.[11] Previously, three major lineages of ST131 were identified that differed mainly based on their fimH alleles: A (mainly fimH41), B (mainly fimH22) and C (mainly fimH30).[19] Clade C has predominated since the 2000s, corresponding with the rapid dissemination of the *bla_CTX-M-15_* allele.[11, 19, 20] There is growing evidence in SSA that ST 131 CTX-M-15 *E. coli* strains are increasing, but there remains a limited number of studies assessing the clonality of *E.coli*, the distribution of ST131 and the presence of *bla_CTX-M-15_* genes using whole genome sequencing.[21–24]

In Malawi, specifically, there has been growing concern about increasing rates of antimicrobial resistance to most empirically used antimicrobials.[25, 26] A recent genomic epidemiology study of 94 *E. coli* isolates collected at Queen Elizabeth Central Hospital, a tertiary care center in southern Malawi, and analyzed by whole genome sequencing (WGS), has shown that ST131 is the most common ST in southern Malawi at 14.9% of isolates sequenced. CTX-M-15 was found in 21.4% of ST131 isolates, but occurred across 11 STs.[27] The purpose of our study is to increase our understanding of the genomic epidemiology of *E. coli* in Malawi by conducting WGS, using Oxford Nanopore Technologies (ONT) sequencing performed in Malawi, of isolates collected at a tertiary care hospital in central Malawi. We use this data to define the clonality, virulence genes and antimicrobial resistance genes in the central region of the country and compare these results to southern Malawi.

## METHODS

### Sample Selection

Sixty isolates were selected from the UNC Project-Malawi archives for sequencing. Samples were collected between 2012 and 2018. Isolates were selected on the basis of diversity of phenotypic resistance pattern and clinical source of isolation, similar to a previous study in Malawi.[27]

### Antimicrobial Resistance Testing

The Kirby-Bauer disk diffusion method was used to measure the *in vitro* susceptibility of bacteria to antimicrobial agents at the UNC Project-Malawi microbiology laboratory at the time of isolate collection. Results were obtained with disk diffusion tests that use the principle of standardized methodology and zone diameter measurements correlated with minimum inhibitory concentrations (MICs) with strains known to be susceptible and resistant to various antibiotics. All aspects of the procedure were standardized as recommended by the Clinical and Laboratory Standards Institute (CLSI) in the document “ *Performance Standards for Antimicrobial Susceptibility Testing”*.[28]

### Whole genome sequencing

Overnight cultures of the isolates were grown in 5ml of LB broth at 37°C. Cell pellets from the broth culture were recovered from 1.5ml centrifuged at 10,000g for 2 minutes. Cell pellets were resuspended in 100ul of nuclease-free water. DNA was extracted from the resuspended pellet using the Zymo Quick-DNA Microprep Kit (cell suspensions protocol) per manufacturer’s instructions (Zymo Research, Irvine, CA, USA). The DNA was quantified using the dsDNA kit on a Qubit 2.0 (Thermo Fisher, Waltham, MA, USA). Equal amounts of DNA (~100ng) from each isolate were used for library preparation using the Rapid Barcoding (RBK-004) per manufacturer’s protocol (Oxford Nanopore Technologies, Oxford, UK). Pooled libraries of 12 isolates were run on a R9.4.1 flow cell on a MinION/MinIT for 24 hours at UNC Project-Malawi. Each flow cell was washed once per protocol and a second set of 12 isolates were run for an additional 24 hours or until all pores were exhausted.

### *De novo* genome assembly

Base calling of fast5 files was done with `Guppy` (version 3.4.5) with the R9.4.1 “high accuracy” model.[29] Samples were demultiplexed using a custom tool, `depore` (version 0.1; https://github.com/txje/depore). Reads for each isolate were *de novo* assembled using `Flye` (version 2.7).[30] Assemblies were then polished four times with `racon` (version 1.3.2) and underwent a final polish with `medaka` (version 0.6.2)(http://github.com/nanoporetech/medaka).[31, 32]

### Typing of isolates

Each assembly was aligned using `minimap2` to databases of known serotypes, genomic and plasmid sequence types including plasmid incompatibility, virulence factors, and antimicrobial resistance genes, listed in **Supplemental Table 1**.[33–38] Matches were made for each assembly using similar cutoffs to those used by the Center for Genomic Epidemiology (CGE; https://cge.cbs.dtu.dk/services/data.php) tools, where at least 60% of the feature must match at >90% sequence identity.

### Species identification

We detected O and/or H serotypes and multi-locus sequence types in 59 of 60 presumed *E. coli* samples. The remaining sample appeared to have assembled well and a BLAST search revealed it was a strain of *Klebsiella*. We used this sequence as an outgroup in our phylogenetic analysis and otherwise excluded it from further analysis. An additional isolate was excluded from analysis for poor sequencing coverage.

### Phylogenomics

Assembled genomes were aligned to a set of single-copy orthologs largely conserved across *Enterobacterales* (BUSCO v4, https://busco.ezlab.org/). A subset of 92 genes were selected that appear in all 58 samples that were included in the final analysis. A multiple-sequence alignment was performed for each gene using `MUSCLE` (3.8.31).[39] `RAxML-ng` was used to generate maximum-likelihood phylogenetics trees for the concatenated alignments with model parameter `‘GTR+G’`.[40] For the comparison to existing Malawi *E. coli* isolates, we used publicly available sequence data shown in **Supplemental Table 2**. For each of these samples, a genome was assembled from Illumina sequence data with `SPAdes` (3.14.0) using default parameters.[41] A phylogeny incorporating these sequences was generated as above, using a subset of 80 genes present in all 149 genomes.

### Data Analysis

Data were analyzed using STATA/SE version 16.1 (STATA Inc, College Station, TX). Descriptive statistics were used to describe clinical characteristics, frequencies of gene detections, and disc diffusion results. Given the small overall sample size, Fisher’s exact tests were used to assess for associations between genes detected and phenotypic resistance testing by Kirby-Bauer disk diffusion

## RESULTS

### Isolate characteristics and sequencing data

Clinical and patient characteristics for the isolates in the final analysis are summarized in **Table 1**. The majority were from a urinary source (59%) and were from females (74%). Blood (24%) was the second most common site. Patient white blood cell counts and hemoglobin levels were available for 20 patients; medians and interquartile ranges are included in **Table 1.** Five sequencing runs generated 9,291,492 reads totalling 40.8 Gbp with a mean read length of 4,391 bp and N50 of 8,732 bp. Sequencing summary statistics across samples are described in **Table 2** for all 60 isolates. Two isolates were excluded from further analysis for poor sequencing coverage (1) or being identified as a different species (1), resulting in 58 isolates included in all analyses. Kirby-Bauer antimicrobial susceptibility testing results are summarized in **Table 3**. Of note, the majority of isolates were resistant to amoxicillin and trimethoprim-sulfamethoxazole (TMP-SMX) and 57% (33/58) were resistant to ceftriaxone. These are commonly prescribed antibiotics in the outpatient (amoxicillin and TMP-SMX) and inpatient (ceftriaxone) settings in Malawi.[42]

**Table 1.** Patient Characteristics of 58 Included Isolates

| Variable | Number of Isolates<br>n (%) |
| --- | --- |
| Sex |  |
| Female | 43 (74) |
| Male | 15 (26) |
| Age* | 34 (20-47) |
| Sample Type |  |
| Blood | 14 (24) |
| Urine | 34 (59) |
| CSF | 2 (3) |
| Body Fluid | 2 (3) |
| Joint | 2 (3) |
| Other | 4 (7) |
| Lab Values# |  |
| White blood cell<br>count (cells/μL) | 7.5 (3.8-9.1) |
| Hemoglobin (g/dL) | 9.8 (8.4-11.6) |

\*Median (IQR); data available for 54 patients
#Median (IQR); data available for 20 patients

**Table 2.** Sequence Data Characteristics of All Sequenced Isolates

| Sequence Data | Value |
| --- | --- |
| Number of reads | 135,216 (21,843 - 539,714) |
| Total bps | 589 Mbp (84 - 2,076 Mbp) |
| Median fragment size sequenced in sample | 2,452 bp (586 - 4,560 bp) |
| N50 assembly | 4.49 Mbp (0.30 - 5.37 Mbp) |
| Number of contigs | 8.5 (1 - 80) |
| Chromosomal mean coverage | 114 (16 - 417) |

**Table 3.** Kirby-Bauer Antimicrobial Resistance Patterns of Included Isolates*

| Drug (Total Tested) | Susceptible, n (%) | Intermediate, n (%) | Resistant, n (%) |
| --- | --- | --- | --- |
| Amox-Clav (56) | 15 (27%) | 9(16%) | 32 (57%) |
| Amoxicillin (58) | 3 (5%) | 1 (2%) | 54 (93%) |
| Cefotaxime (49) | 21 (43%) | 0 (0%) | 28 (57%) |
| Ceftriaxone (58) | 25 (43%) | 0 (0%) | 33 (57%) |
| Chloramphenicol (48) | 29 (60%) | 4 (8%) | 15 (31%) |
| Ciprofloxacin (58) | 21 (36%) | 2 (3%) | 35 (60%) |
| Gentamicin (57) | 30 (53%) | 0 (0%) | 27 (47%) |
| Imipenem (31) | 31 (100%) | 0 (0%) | 0 (0%) |
| Nalidixic Acid (57) | 17 (30%) | 2 (4%) | 38 (67%) |
| TMP-SMX (50) | 4 (8%) | 1 (2%) | 45 (90%) |

\*: Percentage may not equal 100% due to rounding.

### Sequence types, H and O groups

Twenty-one different ST groups were found among 53 isolates. Five isolates could not be assigned to a unique ST group (**Supplemental Table 3**). ST131 was the most common type identified [19/58 isolates (32.8%)], followed by ST410 which was present in 9 of 58 isolates (15.5%). Other ST that occurred in more than one isolate included ST69 (5.2%), ST38 (3.4%), ST617 (3.8%) and ST12 (3.4%). Of the 19 ST131, 18 of them were fimH30 and therefore clade C. One isolate was fimH27 and classified as clade A. The fimH30 was linked to O25 and O16, while the fimH27 strain was O18 but also ST131. The population contained 23 different O groups (**Supplemental Table 4**), with 4 samples being unable to be called, and 19 different H group calls (**Supplemental Table 5**).

### Population structure of *E. coli* in Malawi

A phylogenetic tree of the isolates with previously reported *E. coli* genomes from southern Malawi is shown in **Figure 1**. We were able to generate high quality genomes for 91 of the previously reported isolates.[27] Of note, the ST131 isiolates cluster into a single group and have relatively little genetic distance between them in the tree, suggesting that the expansion of ST131 in Malawi is a recent event.[27]

**Figure 1.**
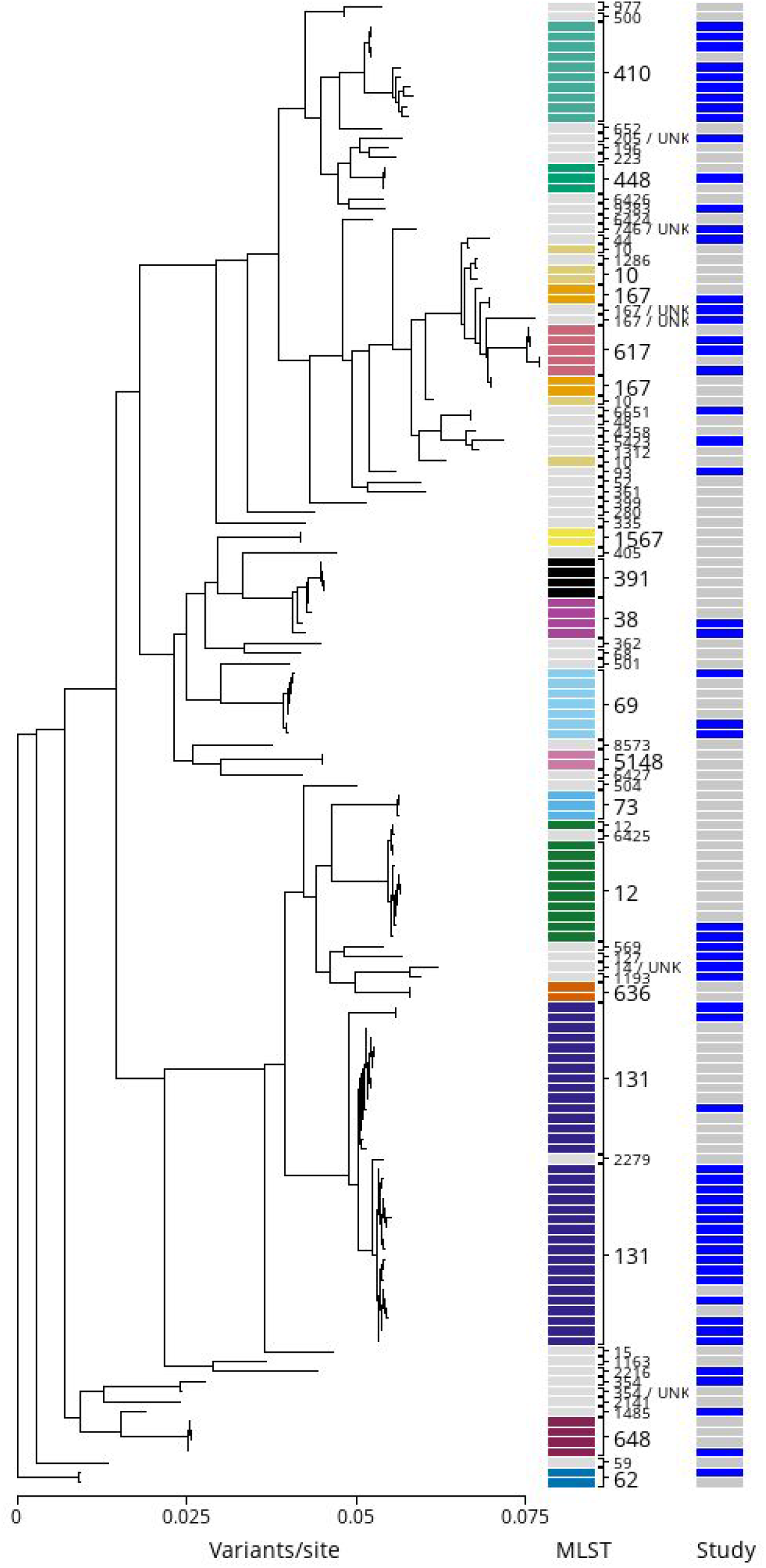
Phylogenetic Tree of Malawian *E. coli* isolates. Phylogenetic relationship among the 58 isolates presented here in relation to previously published Malawi *E. coli* isolates (Supplemental Table 2). Samples cluster by sequence type, but are well-mixed between the two geographically and temporally separated studies. On the right, blue indicates isolates from this study and gray from the previous study.

### Genetic determinants of antimicrobial resistance

Our analysis identified 69 unique AMR genes that are known to encode proteins associated with antimicrobial susceptibility across a range of compounds (**Table 4**). AMR genes occurred across a range of ST and phylogenetic groups (**Figure 2).**

**Figure 2.**
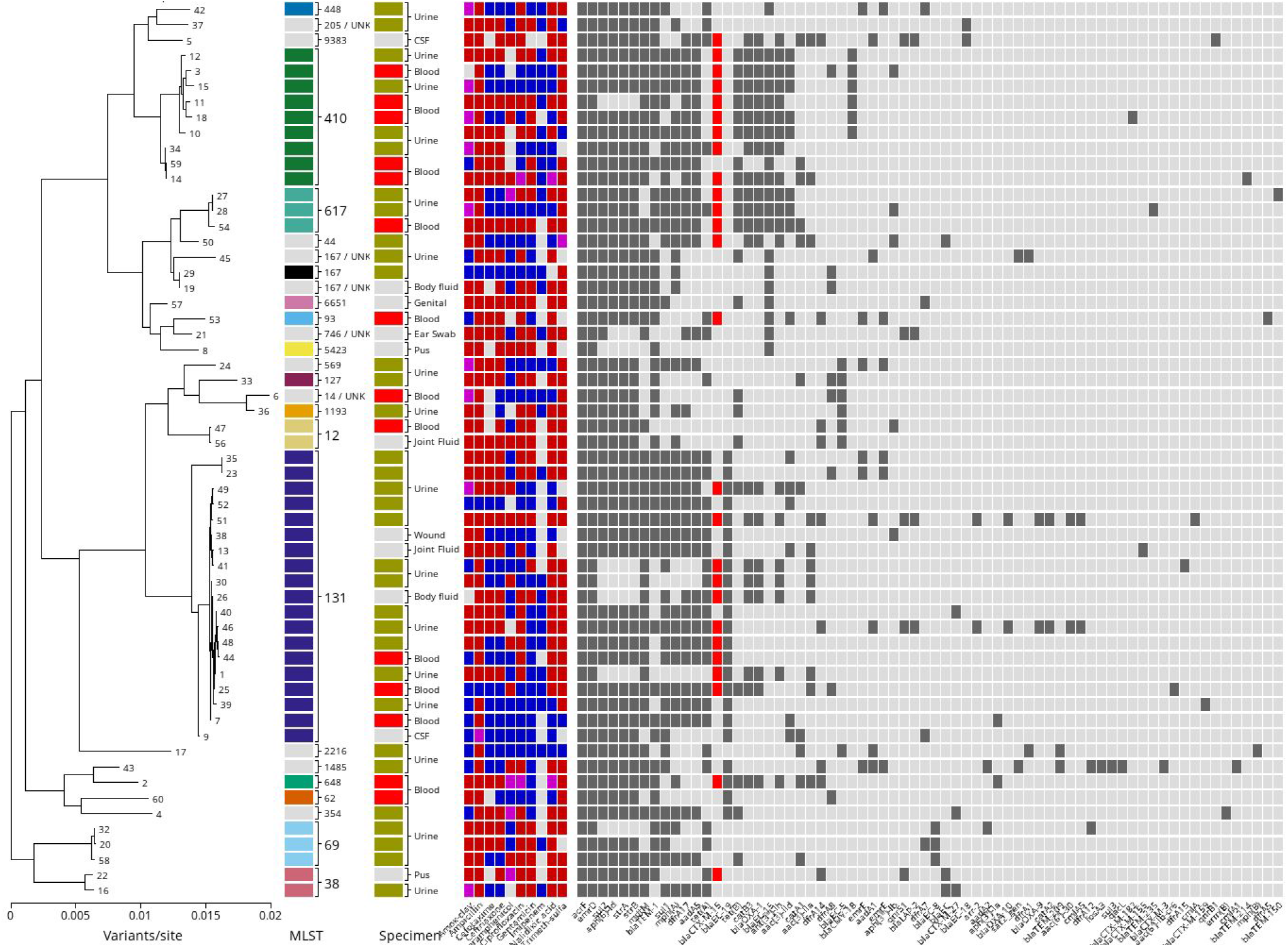
Distribution of Sequence Type, Specimen Type, Resistance Phenotype and Resistance Gene Composition in 58 Malawian *E. coli* Isolates. Phylogenetic relationship among sequenced isolates with corresponding sequence type, specimen type, phenotypic and genomic AMR status. Far left is the phylogeny relating these 58 samples with the sequenced *Klebsiella* from our study as an outgroup (not shown). In line with each terminal branch is the corresponding sample’s sequence type (ST), specimen from which the sample was isolated, AMR phenotype (red: resistant, blue: susceptible, purple: intermediate, gray: unknown), and presence of each detected AMR gene in the genome assembly (dark gray: present, light: absent, red highlights presence of *bla_CTX-M-1_5).*

**Table 4.**
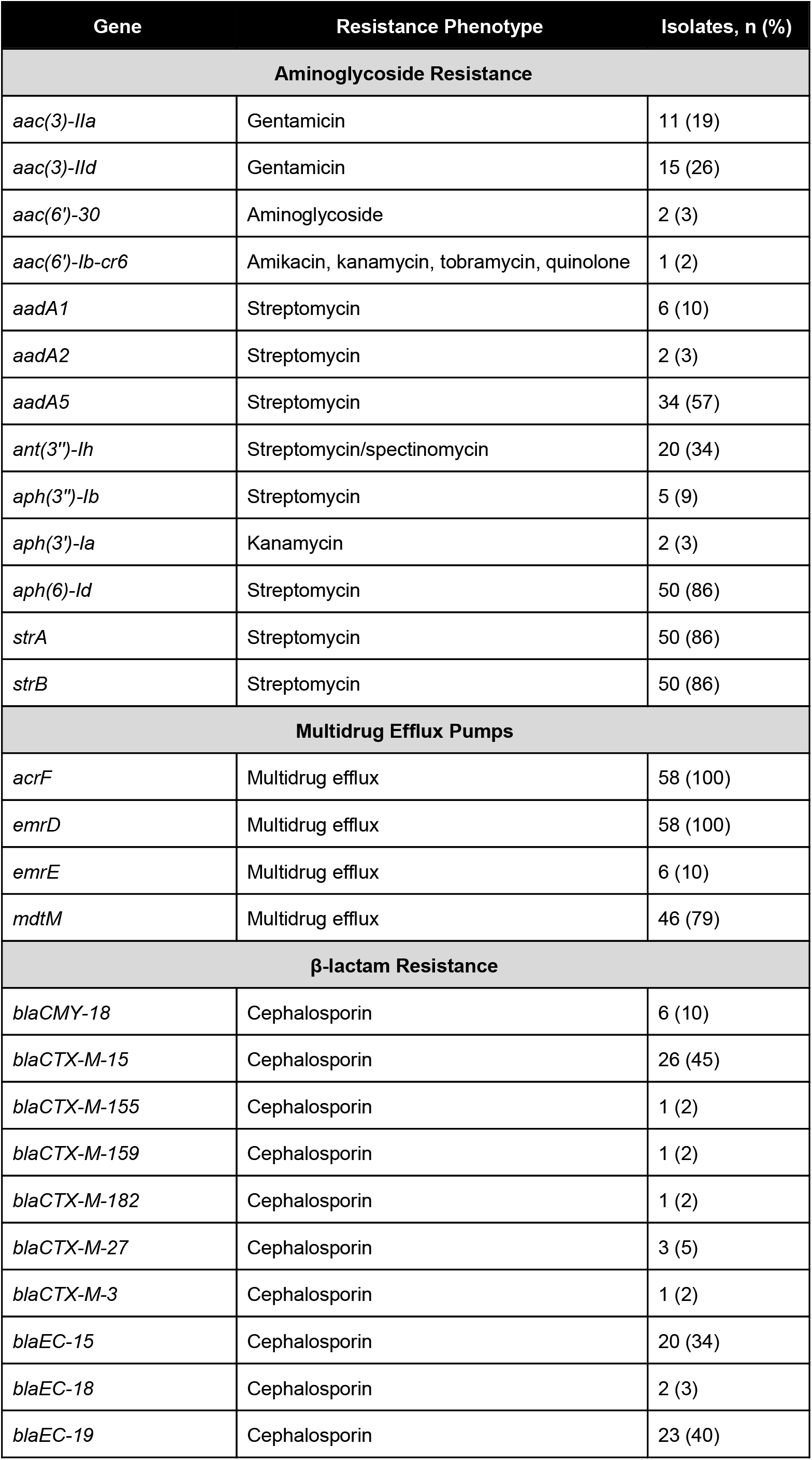
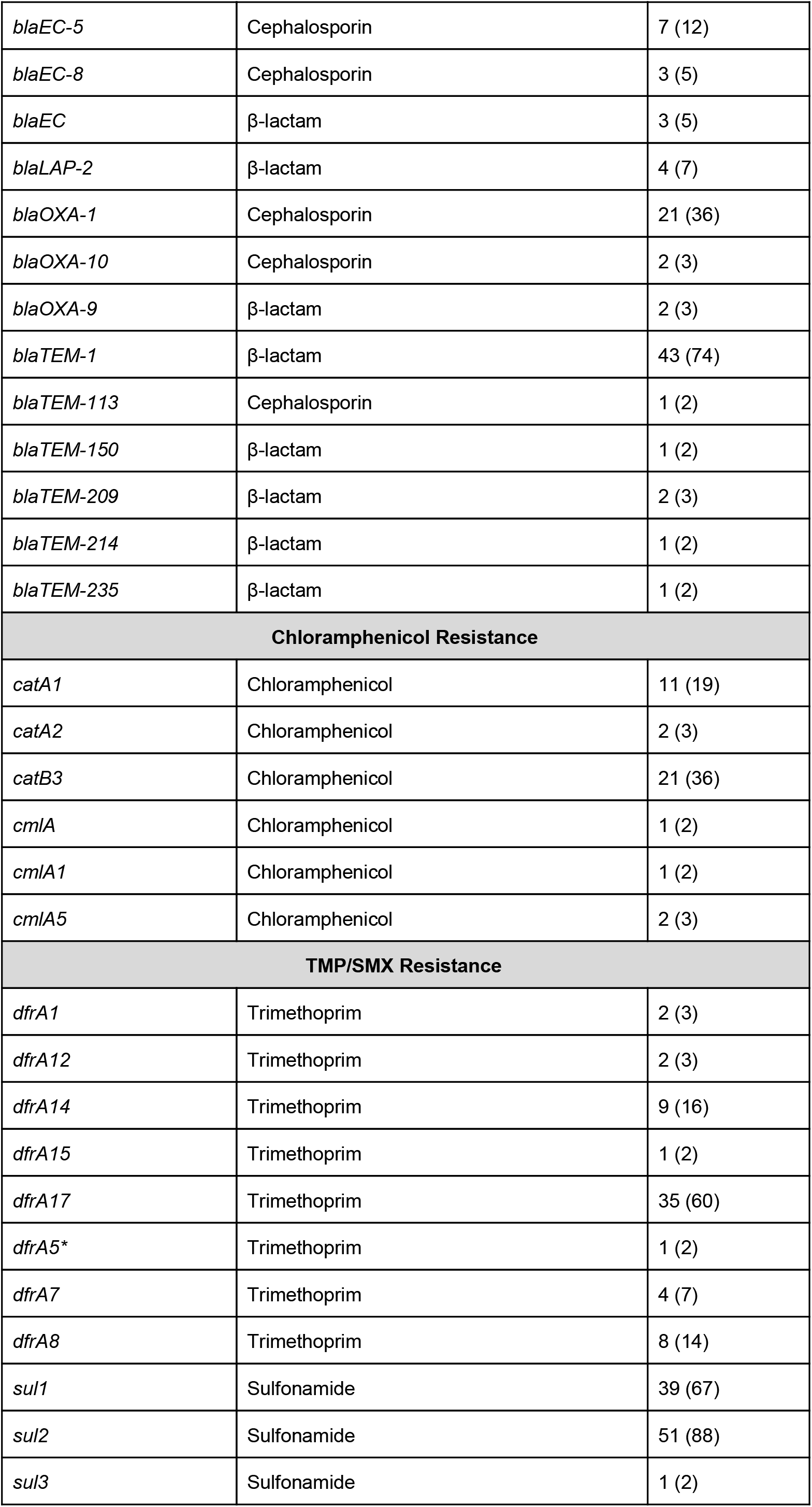
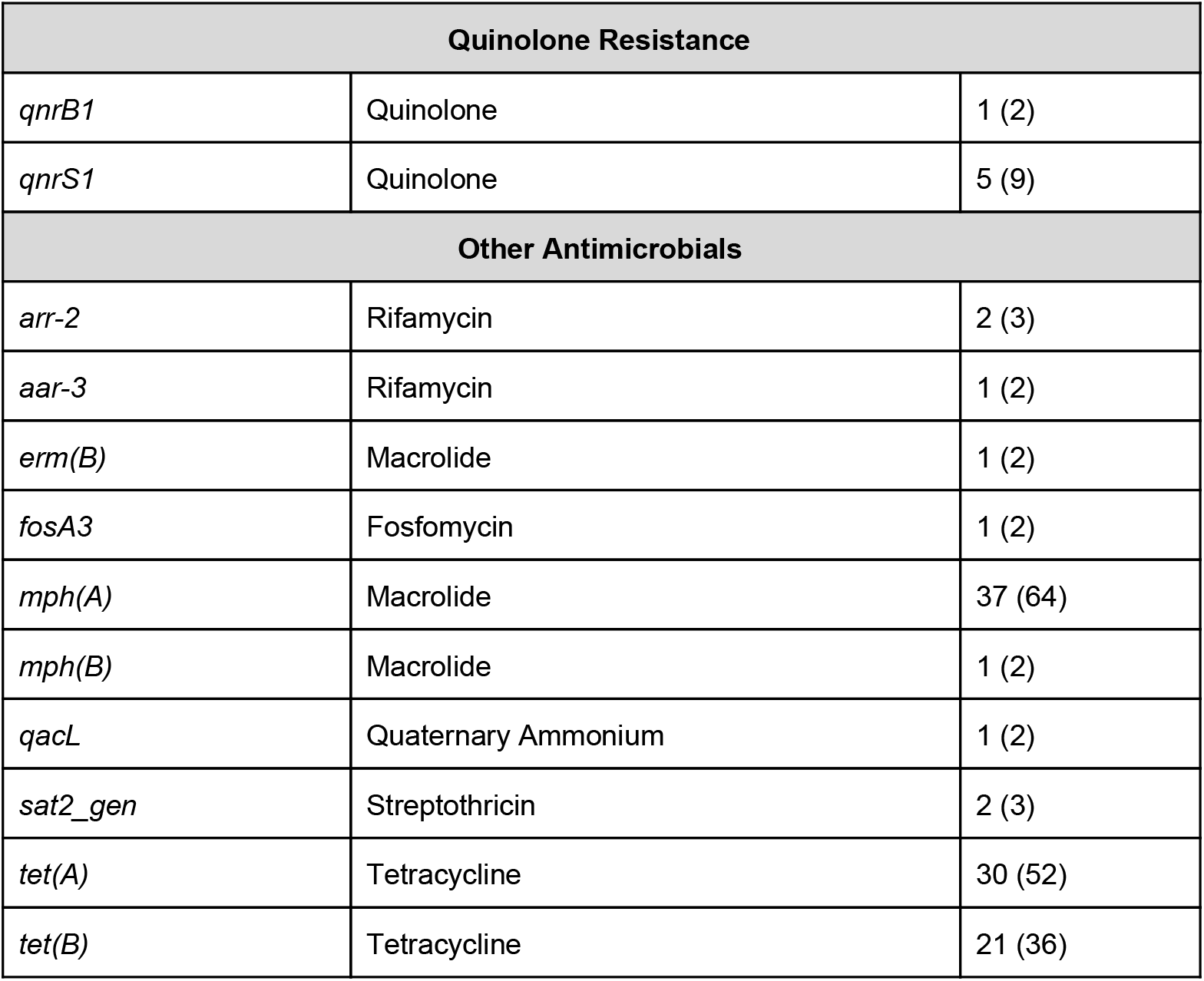
Prevalence of AMR Genes Identified

### β-Lactam and ESBL resistance

We identified 23 genes associated with β-lactam resistance, including 10 ESBL genes (**Table 4**). Extended-spectrum β-lactamases included *bla_TEM-113_, bla_CTX-M-15_, bla_CTX-M-155_, bla_CTX-M-159_, bla_CTX-M-182_, bla_CTX-M-27_, bla_CTX-M3_ bla_EC-15_, bla_EC-18_, bla_EC-19_*. Forty-eight out of 58 (83%) isolates carried at least one ESBL gene, and a total of 40 (69%) isolates carried more than one ESBL gene. The presence of any ESBL gene was associated with phenotypic resistance to ceftriaxone (p<0.001). Fifteen isolates with an ESBL gene retained phenotypic susceptibility to ceftriaxone by disk diffusion. All of these isolates each had only one of the following ESBL genes: *bla_EC-15_, bla_EC-18_, bla_EC-19_*.

The most common ESBL gene detected was *bla_CTX-M-15_*, which occurred in 26 (44.8%) isolates (**Table 5**). Presence of the *bla_CTX-M-15_* gene was associated with ceftriaxone resistance, with 26/26 isolates being resistant to ceftriaxone by disk diffusion. The *bla_CTX-M-15_* gene was found in isolates of 8 different sequence types, most commonly ST131 [10/19 isolates (52.6%)] and ST410 [8/9 (88.9%) isolates]. We identified 15 O25b:H4-ST131 isolates, of which 10 (66%) had the *bla_CTX-M-15_*gene. In ST131, the *bla_CTX-M-15_*gene was highly associated with a chromosomal location with 8 isolates (80%) having chromosomal copies, one isolate had a plasmid copy and one isolate with the gene in both locations. ST410 had a very different distribution with 6 of the 8 isolates (75%) containing the *bla_CTX-M-15_* gene in a plasmid mediated copy, while only a single isolate had a chromosomal copy and a single isolate had it in both locations.

**Table 5.** Characteristics of CTX-M-15 Associated Isolates and Ceftriaxone Disk Diffusion Results

| Isolate | Year Collected | Source | Sequence Type (ST) | Plasmid Inclusion Group | Disk Diffusion Result | Location of Gene |
| --- | --- | --- | --- | --- | --- | --- |
| 1 | 2018 | Urine | 131 | inci1 | R | Genomic |
| 2 | 2017 | Blood | 648 | incf | R | Genomic |
| 3 | 2018 | Urine | 410 | incf | R | Plasmid |
| 5 | 2018 | Blood | 9398 | inchi2 | R | Plasmid |
| 10 | 2018 | Body Fluid | 410 | incf | R | Plasmid |
| 11 | 2017 | Urine | 410 | incf | R | Plasmid |
| 12 | 2017 | Urine | 410 | incf | R | Both |
| 14 | 2014 | Pus | 410 | incf | R | Plasmid |
| 15 | 2018 | Urine | 410 | incf | R | Plasmid |
| 18 | 2018 | Body Fluid | 410 | incf | R | Genomic |
| 22 | 2018 | Blood | 38 | incf/incn | R | Plasmid |
| 25 | 2014 | Urine | 131 | incf | R | Genomic |
| 26 | 2014 | Urine | 131 | incf | R | Plasmid |
| 27 | 2015 | Urine | 617 | incf | R | Both |
| 28 | 2014 | Urine | 617 | incf/incn | R | Plasmid |
| 30 | 2016 | Urine | 131 | incf | R | Genomic |
| 34 | 2016 | Urine | 410 | incf | R | Plasmid |
| 41 | 2013 | Blood | 131 | incn | R | Genomic |
| 44 | 2013 | Urine | 131 | incf | R | Genomic |
| 46 | 2013 | Urine | 131 | incf | R | Genomic |
| 48 | 2014 | Blood | 131 | incf | R | Genomic |
| 49 | 2013 | Blood | 131 | incf | R | Genomic |
| 50 | 2013 | Joint Fluid | 44 | incf | R | Plasmid |
| 51 | 2013 | Genital Swab | 131 | incf | R | Both |
| 53 | 2015 | Blood | 93 | inci1/incf | R | Plasmid |
| 54 | 2013 | CSF | 617 | incf | R | Plasmid |

ND: Not Determined; CSF: cerebrospinal fluid

### Fluoroquinolone resistance

We extracted the sequence of the *gyrA* gene from each isolate and translated them into protein sequences. We evaluated for the presence of *gyrA* mutations that have previously been described in Malawi[27]. We identified the S83L mutation in 24 isolates and the D87N mutation in 22 isolates. In addition, we identified the *qnrB1* gene and the *qnrS1* gene in 1 (2%) and 5 (9%) of isolates, respectively. These genes did not co-occur in isolates with *gyr* mutations.

### Aminoglycoside resistance

We identified 13 genes associated with aminoglycoside resistance in these isolates (**Table 4**). Several genes were associated with gentamicin resistance by disk diffusion - *aac(3)-IIa* (p<0.001), *aac(3)-IId* (p<0.001), and *ant(3)-Ih* (p<0.001). We also detected *strA* or *strB* in combination in 50 of the isolates, as has been seen in southern Malawi previously.[27] When found together, these genes confer resistance to streptomycin, which is used in tuberculosis therapy in Africa.[43]

### Resistance to other antimicrobials

We identified 6 genes associated with chloramphenicol resistance, the most common being *catA1* in 11 (19%) of isolates and *catB3* in 21 (36%) of isolates. We did not identify isolates with *floR*, which has previously been identified in southern Malawi [27]. Detection of *catA1* was associated with intermediate susceptibility or resistance to chloramphenicol by disk diffusion (p<0.001), but detection of *catB3* was not (p=0.592). We identified 11 genes associated with resistance to TMP-SMX, the most common being the trimethoprim resistance gene *dfrA17* (60% of isolates) and the sulfonamide resistance genes *sul1* (67% of isolates) and *sul2* (88% of isolates). Of these three genes, only *sul2* detection was associated with TMP-SMX intermediate susceptibility or resistance by disk diffusion (p=0.007); all isolates that were phenotypically intermediate or resistant to TMP-SMX had *sul1* and/or *sul2*. We detected *sul1* and *sul2* in 35 of the 58 isolates. Finally, we identified a handful of resistance genes to other antimicrobials, including rifamycin, macrolides, fosfomycin, and tetracyclines (**Table 4**). Interestingly, the most common AMR genes detected were multidrug efflux pumps, *acrF* and *emrD*, both of which occurred in all isolates.

### Plasmid incompatibility group

We identified a total of 7 different combinations of plasmid incompatibility groups (**Table 6**). The most common was the incf incompatibility group, which was found in nearly 73% of isolates, and although the majority of ST131 were this group, it was not associated with ST131 (p=0.521).

**Table 6.** Prevalence of Plasmid Incompatibility Groups

| Plasmid Incompatibility Group | Number of Isolates, n (%) |
| --- | --- |
| incf | 42 (72.4%) |
| incf/inchi2 | 1 (1.7%) |
| incf/inci1 | 2 (3.4%) |
| incf/incn | 2 (3.4%) |
| inchi2 | 1 (1.7%) |
| inci1 | 3 (5.2%) |
| incn | 1 (1.7%) |
| ND | 6 (10.3%) |

ND: Not detected

### Virulence genes

In total, we identified 38 virulence genes (Table 7) with each isolate carrying a median of 7 genes (IQR 3-10). The virulence factors spanned a range of functions including acid resistance, adhesion, invasins, metalloproteases, and toxins. The most common gene detected was *gad,* which occurred in all 58 isolates and encodes glutamate decarboxylase, an enzyme linked with bacterial ability to resist environmental stresses.[44] Multiple adhesin proteins were also identified. The most common of these were *papC, papH, papG-II*, and *IpfA*. *papC* (36% of isolates), *papH* (35% of isolates), *papG-II* (35% of isolates) are all involved in pili function. *IpfA* encodes the long polar fimbriae associated with human diarrheal disease, which occurred in 19 of 58 isolates.[45] A single protectin encoded by *iss* was identified and was the second most common virulence factor identified in 35 of 58 isolates. The *iss* (increased serum survival) gene was first identified in a human septicemic *E. coli* isolate and was associated with a 20-fold increase in complement resistance and a 100-fold increase in virulence toward 1-day-old chicks. [46–48] Multiple siderophores were identified, with the most common being *iha*, occurring in 28 of 58 isolates.[49] We also identified two common toxins, *sat* which occurred in 22 of 58 isolates and *senB* in 21 of 58 isolates. The secreted autotransporter toxin *(sat)* appears to fall within one subgroup of autotransporters recently classified as the SPATE (serine protease autotransporters of *Enterobacteriaceae)* family. It acts as a vacuolating cytotoxin for bladder and kidney epithelial cells.[50] *senB* encodes the TieB protein, which may have some role in enterotoxicity of EIEC.[51, 52]

**Table 7.** Prevalence of Virulence Genes Identified

| Category | Gene | Isolate, n (%) |
| --- | --- | --- |
| Acid resistance | *gad* | 58 (100%) |
| Adhesin | *afaC* | 13 (22%) |
|  | *air* | 9 (16%) |
|  | *lpfA* | 19 (33%) |
|  | *nfaE* | 13 (22%) |
|  | *papA* | 1 (2%) |
|  | *papC* | 21 (36%) |
|  | *papE* | 2 (3%) |
|  | *papG-II* | 20 (35%) |
|  | *papG-III* | 1 (2%) |
|  | *papH* | 20 (35%) |
|  | *sfaF* | 3 (5%) |
|  | *sfaS* | 1 (2%) |
| Effector (T3SS) | *espI* | 1 (2%) |
| Invasin | *ibeA* | 1 (2%) |
| Metalloprotease | *sslE* | 23 (40%) |
| Microcin | *mchB* | 2 (3%) |
|  | *mchC* | 3 (5%) |
|  | *mchF* | 6 (10%) |
|  | *mcmA* | 3 (5%) |
| Protectin | *iss* | 35 (60%) |
| Regulator | *eilA* | 9 (16%) |
| Siderophore | *iha* | 29 (50%) |
|  | *ireA* | 6 (10%) |
|  | *iroN* | 8 (13%) |
|  | *iutA* | 3 (5%) |
| Toxin | *cma* | 1 (2%) |
|  | *cnf1* | 6 (10%) |
|  | *hly-alpha* | 9 (16%) |
|  | *pic* | 1 (2%) |
|  | *sat* | 23 (40%) |
|  | *senB* | 22 (37%) |
|  | *tsh* | 1 (2%) |
|  | *vat* | 6 (10%) |
| **Transferase** | *capU* | 7 (12%) |
| **ATP-binding cassette transporter** | *ybtP* | 20 (35%) |
|  | *ybtQ* | 22 (38%) |

## DISCUSSION

In this study, we sequenced 58 *E. coli* isolates collected at a tertiary care center in Lilongwe, Malawi between 2012-2018. To our knowledge, this is one of the first studies from Malawi demonstrating the feasibility of performing all steps of the whole genome sequencing process, including DNA extraction, library preparation, and the sequencing itself, on site in a local laboratory.

Relatively little data concerning the genomics of *E. coli* exist in Africa to date.[21–24] The isolates sequenced as part of our project collected in Lilongwe, in the central region of Malawi, are highly phylogenetically similar to those seen in a previous study conducted in Blantyre, Malawi, in the southern region.[27] Interestingly, there are multiple branches that contain nearly identical isolates sequenced between the two studies despite the fact that the collection sites are over 300km apart from each other and our isolates were on average collected several years later. Similar to the previous report, we identified a diverse set of AMR genes, with similar genes occurring across a range of *E. coli* lineages in Malawi. We also provide additional information on common virulence genes associated with *E. coli* in Malawi, many of which are shared across a range of lineages.

Consistent with the Blantyre study’s findings, we show that *E. coli* is highly diverse, with a distribution of ST similar to global isolates. In our collection, ST131 isolates are within a clade with relatively little genetic distance within it, suggesting a recent expansion of this ST in Malawi. Importantly, the previous study utilized samples collected between 2006 and 2013, allowing temporal comparisons between studies. Our study suggests that ST 131 has become more prevalent (14.9% increasing to 32.8%) and that the *bla_CTX-M-15_* gene is occurring at a higher frequency (21.3% increasing to 44.8%) in the intervening years. This is consistent with global trends that suggest that the highly drug-resistant *E. coli* ST131, associated with the *bla_CTX-M-15_* gene, has been increasing in prevalence.[9–11] [15–18] There is some subtle structure within ST131, with isolates from this study that are primarily localized on a branch with a longer internal branch length, which may suggest the seeding and expansion locally in central Malawi (**Figure 1**). Another difference from previous work is the higher number of ST410 isolates in this study, which carried the *bla_CTX-M-15_* gene at the highest frequency of all the sequence types. Overall, there does not appear to be any strong associations between overall pattern of AMR gene content, virulence gene content, isolation site, and ST within the isolates (**Figure 2**), similar to previous reports.[27]

Although globally there is a strong association between ST131 and presence of the *bla_CTX-M-15_* gene, here we identify the *bla_CTX-M-15_* gene across a diverse set of lineages.[11] This finding is similar to other studies in Tanzania and Malawi, where the *bla_CTX-M-15_* gene was found across numerous ST.[22, 27] We see the *bla_CTX-M-15_* gene in 8 different sequence types, including ST131. The ST that most commonly contained the *bla_CTX-M-15_* gene was ST410. All of the ST131 isolates that contained the *bla_CTX-M-15_* gene were O25b:H4-ST131 in this study. Overwhelmingly, the ST131 isolates were clade C, containing the fimH30 gene, consistent with the global expansion of ST131-H30.[12] Interestingly, there is a strong association between ST and where the *bla_CTX-M-15_* gene is carried in the ST131 and ST410 isolates. ST131 was much more likely to have a genomic location for the gene, whereas ST410 more frequently carried the gene on a plasmid. This may have implications for how the *bla_CTX-M-15_* gene spreads in Malawi. Given the diverse lineages that carry the *bla_CTX-M-15_* gene, additional studies are needed to better understand the epidemiology of this gene in Malawi.

Overall, the patterns of AMR gene prevalence were similar to the one previous report from southern Malawi. *sul2*, *strA*, *strB*, *dfrA*, *bla_TEM-1_*, and *sul1* genes remained very common in the population. Interestingly, chloramphenicol resistance gene prevalence was lower than previously reported, with a decrease in the prevalence of the types of *catA* 64.9% to 22%.[27] This potentially reflects decrease use of the drug in the community with changing treatment guidelines and increasing availability of alternative agents with fewer side effects.[42] Most resistance genes were detected broadly across different genotypes in this study (**Figure 2**).

In summary, we confirm that the *E. coli* population in Malawi is highly diverse, with evidence for the ongoing recent expansion of the ST131 group in the country. We see a higher proportion of ST 131 isolates and a higher prevalence of the *bla_CTX-M-15_* gene in our isolates, which were collected a few years later than previous reports. This expansion is consistent with the global increase in O25b:H4-ST131 bearing the fimH30 gene. *E. coli* genotypes are similar between two major tertiary care hospitals that are quite distant, with highly related isolates being found between the sites. As previously seen, AMR genes, including the *bla_CTX-M-15_* gene, are broadly contained across sequence types. A high diversity of virulence genes were seen within *E. coli* isolates. These data were collected by conducting ONT sequencing in Malawi, highlighting the possibility of conducting rapid longitudinal genomic epidemiology studies of consequential bacteria in sub-Saharan Africa where the sequencing is conducted on site.

## AUTHOR STATEMENTS

### Authors and contributors

GT contributed to investigation, data curation, and writing-review and editing. EJC contributed investigation, data curation, formal analysis, writing-original draft preparation, and project administration. RK contributed investigation, resources and writing-review and editing. JK, TC, and IN contributed investigation and data curation. DvD contributed writing-review and editing. IH contributed resources, writing-review and editing, and conceptualization. JJJ contributed conceptualization, methodology, formal analysis, resources, data curation, writing-original draft preparation, supervision, project administration, and funding. JW contributed conceptualization, methodology, software, formal analysis, resources, data curation, writing-original draft preparation, supervision, visualization, and project administration.

### Ethical approval

Secondary use of stored clinical isolates from UNC Project Malawi was approved by the UNC IRB (18-3292) and National Health Sciences Research Committee IRB (approval number 2205). This work was deemed non-human subjects research.

## Acknowledgements

We would like to thank the laboratory team and management at UNC Project-Malawi for hosting the project. Thank you to the unknown individuals for use of their isolates.

