## Supplemental Material for "Genomic Epidemiology of *Escherichia coli* Isolates from a Tertiary Referral Center in Lilongwe, Malawi"

### SUPPLEMENTARY MATERIAL

#### Supplemental Table 1. Databases Used for Sequence Analysis

| Database | Version/Date | Ref |
|---|---|---|
| MLST | https://bitbucket.org/genomicepidemiology/mlst_db/commits/94abfd0 (2020-03-10) | Center for Genomic Epidemiology |
| pMLST | https://bitbucket.org/genomicepidemiology/pmlst_db/commits/a100502 (2020-03-01) | Center for Genomic Epidemiology |
| VirulenceFinder | https://bitbucket.org/genomicepidemiology/virulencefinder_db/commits/13d72a8 (2020-03-01) | Center for Genomic Epidemiology |
| fimTyper (fimH) | https://bitbucket.org/genomicepidemiology/fimtyper_db/commits/f999a42 (2020-03-10) | Center for Genomic Epidemiology |
| SRST2 (serotype) | https://github.com/katholt/srst2/commit/fe027e55 (2020-03-01) | http://genomemedicine.com/content/6/11/90 |
| AMRFinderPlus | 3.6 (2020-01-06.1) | NCBI AMR Database (https://www.ncbi.nlm.nih.gov/bioproject/PRJNA313047) |

### Supplemental Table 2: Public Sequences Used in Analysis

| Source | Strain ID | ENA Accession ID |
|---|---|---|
| Musicha et al. [30] | A39011 | ERS668966 |
| Musicha et al. [30] | 522_A | ERS668975 |
| Musicha et al. [30] | A45214 | ERS668976 |
| Musicha et al. [30] | 1010805 | ERS668977 |
| Musicha et al. [30] | BHA15G | ERS668978 |
| Musicha et al. [30] | BKQ7M8 | ERS668979 |
| Musicha et al. [30] | BKR1Z7 | ERS668980 |
| Musicha et al. [30] | D40034 | ERS668981 |
| Musicha et al. [30] | BKQ5JN | ERS668982 |
| Musicha et al. [30] | A7898 | ERS668983 |
| Musicha et al. [30] | 3361 | ERS668984 |
| Musicha et al. [30] | D3787 | ERS668967 |
| Musicha et al. [30] | BKR406 | ERS668987 |
| Musicha et al. [30] | D3475 | ERS668988 |
| Musicha et al. [30] | B12381 | ERS668989 |
| Musicha et al. [30] | A7503 | ERS668990 |
| Musicha et al. [30] | C301 | ERS668991 |
| Musicha et al. [30] | A38084 | ERS668992 |
| Musicha et al. [30] | D3275 | ERS668993 |
| Musicha et al. [30] | A5175 | ERS668994 |
| Musicha et al. [30] | B1PG3 | ERS668968 |
| Musicha et al. [30] | D4531 | ERS668995 |
| Musicha et al. [30] | 2473 | ERS668996 |
| Musicha et al. [30] | D3420 | ERS668998 |
| Musicha et al. [30] | C10382 | ERS668999 |
| Musicha et al. [30] | 2228 | ERS669000 |
| Musicha et al. [30] | 10151 | ERS669001 |
| Musicha et al. [30] | 8728 | ERS669002 |
| Musicha et al. [30] | 10129 | ERS669004 |
| Musicha et al. [30] | A36329 | ERS669005 |

| | | |
|---|---|---|
| Musicha et al. [30] | C1289 | ERS668969 |
| Musicha et al. [30] | B9070 | ERS669006 |
| Musicha et al. [30] | 9597 | ERS669007 |
| Musicha et al. [30] | D36115 | ERS669008 |
| Musicha et al. [30] | 2209 | ERS669009 |
| Musicha et al. [30] | A25576 | ERS669010 |
| Musicha et al. [30] | D37334 | ERS669011 |
| Musicha et al. [30] | D25640 | ERS669012 |
| Musicha et al. [30] | D25641 | ERS669013 |
| Musicha et al. [30] | D29454 | ERS669014 |
| Musicha et al. [30] | A38988 | ERS669015 |
| Musicha et al. [30] | 1012184 | ERS668970 |
| Musicha et al. [30] | A40286 | ERS669016 |
| Musicha et al. [30] | 4464 | ERS669017 |
| Musicha et al. [30] | B9222 | ERS669018 |
| Musicha et al. [30] | C14036 | ERS669019 |
| Musicha et al. [30] | A36140 | ERS669020 |
| Musicha et al. [30] | 3524 | ERS669021 |
| Musicha et al. [30] | 4600 | ERS669022 |
| Musicha et al. [30] | C12359 | ERS669024 |
| Musicha et al. [30] | A32883 | ERS669025 |
| Musicha et al. [30] | BKQA8N | ERS668971 |
| Musicha et al. [30] | 9693 | ERS669027 |
| Musicha et al. [30] | 2558 | ERS669028 |
| Musicha et al. [30] | A27 | ERS669029 |
| Musicha et al. [30] | C29 | ERS669031 |
| Musicha et al. [30] | A333 | ERS669032 |
| Musicha et al. [30] | D39719 | ERS669033 |
| Musicha et al. [30] | C30 | ERS669034 |
| Musicha et al. [30] | C15 | ERS669035 |
| Musicha et al. [30] | D40059A | ERS669036 |
| Musicha et al. [30] | BHAIAI | ERS668972 |

| | | |
|---|---|---|
| Musicha et al. [30] | D43713 | ERS669037 |
| Musicha et al. [30] | C4 | ERS669039 |
| Musicha et al. [30] | A16 | ERS669041 |
| Musicha et al. [30] | A35440 | ERS669042 |
| Musicha et al. [30] | C33B | ERS669044 |
| Musicha et al. [30] | D32322 | ERS669045 |
| Musicha et al. [30] | C14 | ERS669047 |
| Musicha et al. [30] | D3871 | ERS669048 |
| Musicha et al. [30] | 1014142 | ERS669049 |
| Musicha et al. [30] | D29253 | ERS668973 |
| Musicha et al. [30] | B28 | ERS669050 |
| Musicha et al. [30] | C20b | ERS669051 |
| Musicha et al. [30] | D49086 | ERS669052 |
| Musicha et al. [30] | D48799 | ERS669054 |
| Musicha et al. [30] | B3 | ERS669055 |
| Musicha et al. [30] | 1016948 | ERS668974 |
| Musicha et al. [30] | BKQ79K_1 | ERS669067 |
| Musicha et al. [30] | 10276 | ERS669087 |
| Musicha et al. [30] | D26076 | ERS669089 |
| Musicha et al. [30] | A7881 | ERS669101 |
| Musicha et al. [30] | D45621 | ERS669107 |
| Musicha et al. [30] | D4275 | ERS669114 |
| Musicha et al. [30] | A44893 | ERS669128 |
| Musicha et al. [30] | A1a | ERS669146 |
| Musicha et al. [30] | 10140 | ERS668997 |
| Musicha et al. [30] | D42544 | ERS669030 |
| Musicha et al. [30] | D46760 | ERS669038 |
| Musicha et al. [30] | A48349 | ERS669040 |
| Musicha et al. [30] | A45016 | ERS669046 |
| Musicha et al. [30] | A3b | ERS669056 |
| Musicha et al. [30] | D33237 | ERS669057 |

**Supplemental Table 3: Distribution of Sequence Types (ST) in 58 Included Isolates**

| Sequence Type (ST) | Number | Frequency |
|---|---|---|
| 131 | 19 | 32.8% |
| 410 | 9 | 15.5% |
| 69 | 3 | 5.2% |
| 38 | 2 | 3.4% |
| 617 | 3 | 5.2% |
| 12 | 2 | 3.4% |
| 354 | 1 | 1.7% |
| 5423 | 1 | 1.7% |
| 2216 | 1 | 1.7% |
| 569 | 1 | 1.7% |
| 127 | 1 | 1.7% |
| 1193 | 1 | 1.7% |
| 58 | 1 | 1.7% |
| 62 | 1 | 1.7% |
| 93 | 1 | 1.7% |
| 167 | 1 | 1.7% |
| 9385 | 1 | 1.7% |
| 1485 | 1 | 1.7% |
| 648 | 1 | 1.7% |
| 44 | 1 | 1.7% |
| 6651 | 1 | 1.7% |
| 14/18/1416/1540/1666/1927/6460/9139/9779 | 1 | 1.7% |
| 167/2815/2821/4183/5507 | 1 | 1.7% |
| 167/693/694/1417/2266/2504/3015/4189/4815/ 5018/6892/7611/9622 | 1 | 1.7% |
| 205/341/2539/5296/5960/7303/7955 | 1 | 1.7% |
| 746/1144/2601/6225/6581/7178/8221/9447 | 1 | 1.7% |

### Supplemental Table 4: Detected O Groups in 58 Included Isolates

| O Antigen | Count | Frequency |
|---|---|---|
| O25 | 15 | 25.9% |
| O8 | 6 | 10.3% |
| Onovel32 | 4 | 6.9% |
| O153var1 | 4 | 6.9% |
| ND | 3 | 5.2% |
| O18 | 2 | 3.4% |
| Onovel14 | 2 | 3.4% |
| O9 | 2 | 3.4% |
| O16 | 2 | 3.4% |
| O4 | 2 | 3.4% |
| O7 | 2 | 3.4% |
| O84 | 1 | 1.7% |
| O11 | 1 | 1.7% |
| O86 | 1 | 1.7% |
| O24 | 1 | 1.7% |
| O15 | 1 | 1.7% |
| O134 | 1 | 1.7% |
| O45 | 1 | 1.7% |
| O6 | 1 | 1.7% |
| Onovel1 | 1 | 1.7% |
| O75 | 1 | 1.7% |
| O100 | 1 | 1.7% |
| O29 | 1 | 1.7% |
| O17 | 1 | 1.7% |
| Onovel32/O9 | 1 | 1.7% |

ND: Not Determined

### Supplemental Table 5. Detected H Groups in 58 Included Isolates

| H-Type | Number | Frequency |
|---|---|---|
| H4/H53/H55/H17 | 21 | 36.2% |
| H9/H55/H17 | 11 | 19.0% |
| H10/H55/H17 | 4 | 6.9% |
| H18/H55/H17 | 3 | 5.2% |
| H5/H53/H55/H17 | 2 | 3.4% |
| H5/H55/H54/H17 | 2 | 3.4% |
| H31/H35/H55/H17 | 2 | 3.4% |
| H37/H55/H17 | 1 | 1.7% |
| H12/H55/H17 | 1 | 1.7% |
| H1/H55/H17 | 1 | 1.7% |
| H34/H55/H17 | 1 | 1.7% |
| H42/H36/H55/H54/H17 | 1 | 1.7% |
| H6/H44/H17 | 1 | 1.7% |
| H5/H55/H17 | 1 | 1.7% |
| H16/H3/H54 | 1 | 1.7% |
| H45/H35/H55/H17 | 1 | 1.7% |
| H84/H23/H36/H55/H17 | 1 | 1.7% |
| H21/H47/H55 | 1 | 1.7% |
| H9/H36/H55/H17 | 1 | 1.7% |
